# Unsaturated intercellular vapor pressure is relevant for leaf water heavy isotope enrichment

**DOI:** 10.1101/2023.09.12.557463

**Authors:** Charlotte Angove, Marco M. Lehmann, Matthias Saurer, Yu Tang, Petri Kilpeläinen, Ansgar Kahmen, Pauliina P. Schiestl-Aalto, Olli-Pekka Tikkasalo, Jaana K. Bäck, Katja T. Rinne-Garmston

## Abstract

Leaf intercellular vapor pressure (*e*_i_) can be unsaturated, but its effect on leaf water heavy isotope enrichment (LWE) has not yet been quantified. We evaluated the ecological relevance of unsaturated *e*_i_ for LWE, i.e., for leaf water oxygen-18 and deuterium enrichment, using data from a boreal forest stand and a large-scale dataset. Unsaturated *e*_i_ can firstly affect LWE by directly decreasing *e*_i_ in the Craig Gordon model (Mechanism 1), which leads to an increased influence of atmospheric vapor isotopic enrichment above source water (Δ_*v*_), and a decreased influence of kinetic fractionation by diffusion through the stomata and boundary layer (ε_k_). Unsaturated *e*_i_ can secondly affect LWE by changing ε_k_ (Mechanism 2). To evaluate the effect of Mechanism 1 to LWE, we employed sensitivity tests on LWE model performance using varying measured intercellular relative humidity (RH_cellular_), or RH_cellular_ fitted to observed LWE. To explore the effects of Mechanism 2 to LWE, we modified the calculation of ε_k_ and observed consequences to LWE predictions. Unsaturated *e*_i_ is relevant to LWE by Mechanism 1, since a lowered RH_cellular_ noticeably changed LWE predictions. It clearly improved deuterium predictions and conditionally improved oxygen-18 predictions. Isotope fractionation by Mechanism 2 is unlikely relevant to oxygen-18 and deuterium enrichment. Unsaturated *e*_i_ must now be recognized as a variable that introduces error to heavy isotope enrichment models and reconstructions from organic material, via Mechanism 1. We suggest a correction for unsaturated *e*_i_ for both oxygen-18 and deuterium enrichment using a variable RH_cellular_ calculated from atmospheric relative humidity.

## Introduction

### Background

Leaf intercellular spaces are specialized locations for gaseous exchange of CO_2_ and water between leaves and the atmosphere. This exchange imprints on the stable isotope signal stored in leaf water, by leaf water heavy isotope enrichment (LWE; Table **1**) (Dongmann *et al*., 1974; Farquhar *et al*., 1989; Flanagan and Ehleringer, 1991; Farquhar, Cernusak and Barnes, 2007). Such LWE is merged into the oxygen-18 (δ^18^O) and deuterium (δ^2^H) stable isotope values of long-term plant bioindicators (Gessler *et al*., 2009; Cernusak and Kahmen, 2013; Cueni *et al*., 2021). For instance, tree-ring δ^18^O is a widely applied tool to study environmental variables, such as temperature, precipitation, atmospheric relative humidity (RH_atm_), and weather phenomena, such as drought (Hartl-Meier *et al*., 2014; Treydte *et al*., 2014; Gessler *et al*., 2018). On the other hand, δ^2^H of tree rings might be indicative for carbon metabolic changes (Lehmann *et al*., 2021; Vitali *et al*., 2022). Such differences in the behavior of δ^18^O and δ^2^H are owing to their different molecular masses, different biosynthetic fractionations, and the covariance between RH_atm_ and water vapor δ^18^O that is not reciprocated by δ^2^H (Cernusak *et al*., 2022; Holloway-Phillips *et al*., 2022). Nevertheless, when δ^2^H and δ^18^O variabilities in tree rings are interpreted using a dual isotope approach, their changes in relative abundance can be used to reconstruct paleoclimatic RH_atm_ (Voelker *et al*., 2014; Hepp *et al*., 2017). Leaf water δ^2^H is useful for another widely used climatic proxy, leaf *n*-alkanes, which can be used for ecohydrological reconstructions (Sachse *et al*., 2012).

**Table 1.**
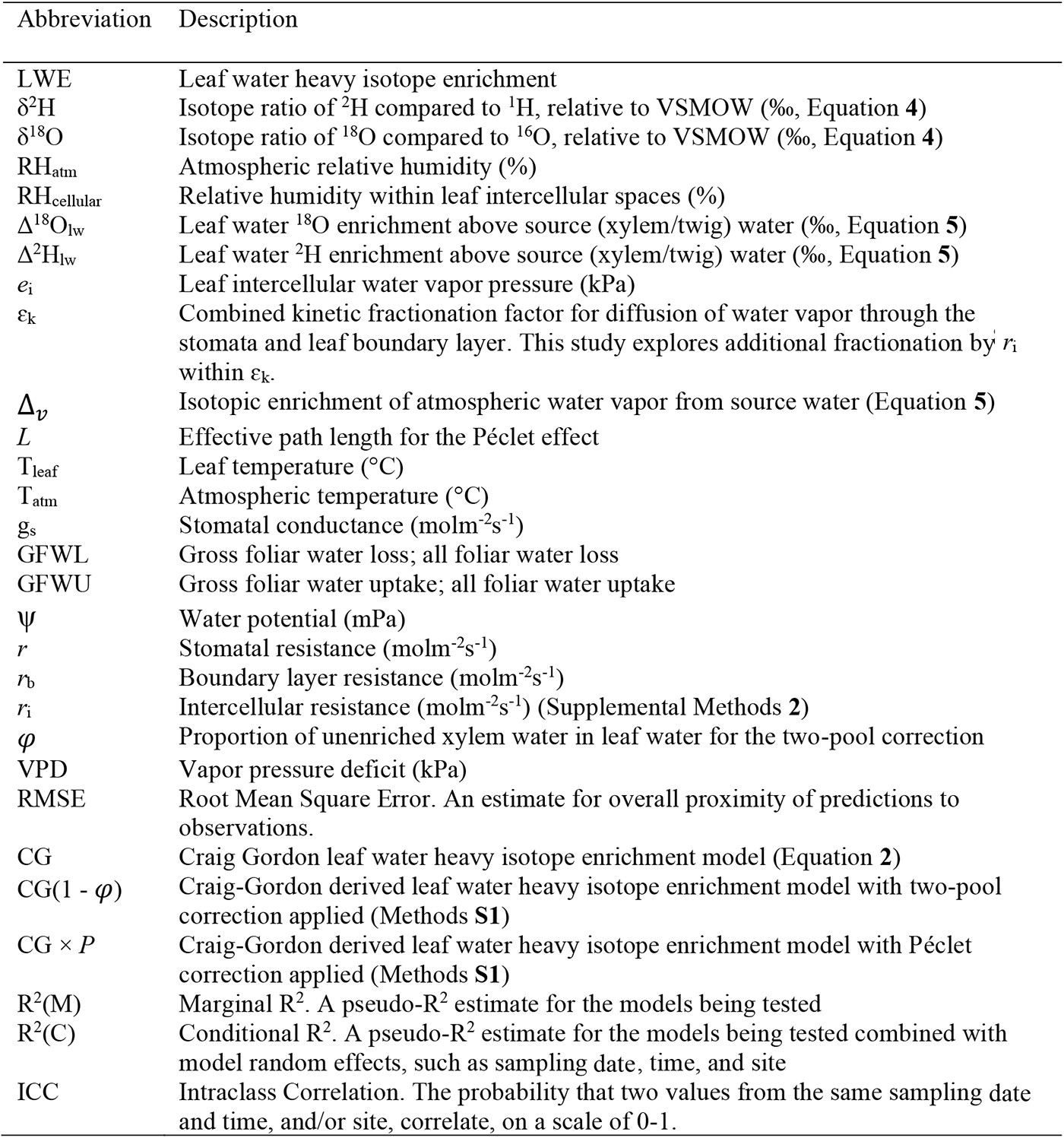
Abbreviations and symbols.

Despite the many climatic and physiological applications of LWE, recent studies challenge our view on the climate related processes regulating LWE, because LWE predictions assume that leaf intercellular vapor pressure (*e*_i_) is saturated, i.e., that RH inside those pores is 100%, while recent studies show that *e*_i_ can be unsaturated, i.e., that RH can drop to as low as 80% (Vesala *et al*., 2017; Cernusak *et al*., 2018; Wong *et al*., 2022). The effect of unsaturated *e*_i_ to LWE is not yet known. If not accounted for, such unsaturated *e*_i_ could be a significant source of error to LWE predictions and reconstructions of past climate and plant response to climate change, via interpretation of tree rings and *n*-alkanes.

Currently, LWE is predicted using an adaptation of a model originally used to predict ocean water heavy isotope enrichment, known as the Craig-Gordon model (Craig, 1965; Dongmann *et al*., 1974; Farquhar *et al*., 1989; Flanagan and Ehleringer, 1991). An approximate calculation for LWE is:

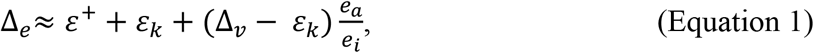

where Δ_*e*_ is the enrichment of the heavy isotope in leaf water above source water, which is Δ^18^O_lw_ for oxygen-18 and Δ^2^H_lw_ for deuterium. Source water is often represented by measured or modelled xylem water isotopic value. Then, ε^+^ is the equilibrium fractionation factor between liquid water and vapor, and ε_*k*_ is the combined kinetic fractionation factor for diffusion of water vapor through the stomata and leaf boundary layer. Next, Δ_*v*_ is the isotopic enrichment of atmospheric water vapor compared to source water, *e*_a_ is the atmospheric water vapor pressure and *e*_i_ is the water vapor pressure in leaf intercellular spaces. Calculations for all variables are demonstrated in supporting information by Cernusak et al. (2022). We will hereafter refer to a more accurately assembled version of Equation **1** (Farquhar, Cernusak and Barnes, 2007):

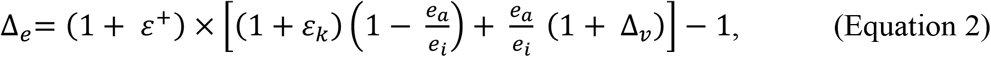

The Craig Gordon model tends to overestimate LWE (Allison, Gat and Leaney, 1985; Leaney *et al*., 1985; Bariac *et al*., 1989; Walker *et al*., 1989). There are three commonly known model corrections to improve LWE model prediction accuracy by considering isotopic inhomogeneities within leaves, and non-steady state conditions. Firstly, the two-pool correction was introduced to reduce LWE overestimation by accounting for the morphological observation that not all leaf water is equally exposed to evaporative enrichment, since most evaporation from leaves occurs at specialized evaporative sites (Leaney *et al*., 1985; Song *et al*., 2015). Then, the Péclet correction was introduced to correct for back-diffusion of heavier stable isotopologues from evaporative sites (Farquhar and Lloyd, 1993). Finally, non-steady state modelling was introduced for circumstances when transpiration rate is low enough that a relatively slow leaf water turnover rate leads to cumulative LWE (Farquhar, Cernusak and Barnes, 2007). But model corrections do not ubiquitously improve LWE predictions across studies, for example, the Péclet correction is unreliable at improving model accuracy for reasons that are not fully understood, related to the effective path-length (*L*), which is more like a “fitting parameter” than a measurable dimension (Cernusak and Kahmen, 2013). Other factors known to affect LWE that have not been accounted for in models include xylem water deuterium inaccuracies by cryogenic water extraction artefacts, and xylem sampling effects (Chen *et al*., 2020; Barbeta *et al*., 2022; Diao *et al*., 2022; Nehemy *et al*., 2022).

## Theory for unsaturated *e*_i_ effects on LWE

### Mechanism 1

Water vapor pressure is saturated when water vapor is in thermodynamic equilibrium with its condensed state. Originally, there were conflicting views about whether leaf intercellular vapor pressure (*e*_i_) is saturated or unsaturated (Jarvis and Slatyer, 1970; Farquhar and Raschke, 1978; Sharkey *et al*., 1982; Canny and Huang, 2006). It was only recently, when the first direct experimental evidence of unsaturated *e*_i_ was released (Cernusak *et al*., 2018; Wong *et al*., 2022). Unsaturated *e*_i_ is particularly relevant to LWE because LWE is caused by the isotopic exchange between leaf water and intercellular vapor. Predictions of LWE rely on an *e*_i_ estimate (Equation **2**). When *e*_i_ is lowered in Equation **2**, it increases the influence of Δ_*v*_, and decreases the influence of ε_*k*_, to LWE by increasing 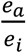 (Mechanism 1, Equation **2**). Since very high RH_atm_ (93%) can affect the influence of atmospheric water vapor isotopologues to LWE (Lehmann *et al*., 2018), an increased influence of Δ_*v*_ by decreased intercellular RH (RH_cellular_) is likely relevant to LWE. Similarly, since ε_*k*_ is renowned to be important for LWE (Farquhar *et al*., 1989), a reduced influence of ε_*k*_will likely have a noticeable impact to LWE. Therefore, the effect of unsaturated *e*_i_ to LWE by Mechanism 1 is likely impactful to LWE.

If *e*_i_ is saturated, it is possible to calculate *e*_i_ using only leaf temperature (T_leaf_; Nobel (2005)). But, since *e*_i_ can be unsaturated, there are more factors that contribute to *e*_i_ than T_leaf_ (Vesala *et al*., 2017; Buckley and Sack, 2019). For instance, changes in leaf-atmosphere water fluxes could interact with *e*_i_. Indeed, unsaturated *e*_i_ could lead to reduced gross foliar water loss (GFWL), if the equation for transpiration, below, can be used to infer effects of unsaturated *e*_i_:

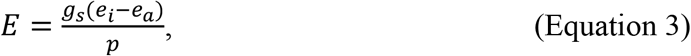

where *E* is transpiration rate, g_s_ is stomatal conductance, and p is air pressure (Farquhar *et al*., 1980). But the effects of unsaturated *e*_i_ can unlikely be evaluated using Equation **3**, because unsaturated *e*_i_ may increase g_s_ (Buckley and Sack, 2019). Indeed, the water potential (ψ) of intercellular spaces is lowered by unsaturated *e*_i_, which arises many questions about our understanding of leaf water transport biology (Buckley and Sack, 2019). Leaf intercellular spaces might withstand lower ψ by unsaturated *e*_i_, via humidity gradients inside of leaf air spaces that reduce ψ differences between leaf cell walls and intercellular spaces, and by concavely curved water-air interfaces in intercellular spaces (Vesala *et al*., 2017; Cernusak *et al*., 2018; Wong *et al*., 2022). Another consequence of unsaturated *e*_i_ is that more water vapor molecules could be up taken from the atmosphere into leaf intercellular spaces, otherwise known as increased gross foliar water uptake (GFWU) which would also depend on RH_atm_ and stomatal conductance (Vesala *et al*., 2017). Overall, there is no empirical evidence showing that unsaturated *e*_i_ would lead to reduced GFWL (Equation **3**) or increased GFWU. Nevertheless, the outcome of both, either reduced GFWL or increased GFWU, contribute to a reduced GFWL:GFWU ratio. The reduced GFWL:GFWU ratio could partly explain the response observed in Equation **2** when *e*_i_ is lowered from saturated vapor pressure, which is currently used by literature, to unsaturated vapor pressure.

### Mechanism 2

When *e*_i_ is unsaturated, it can influence LWE, not only by directly changing *e*_i_ in Equation 2 (Mechanism 1), but also by changing the kinetic fractionation factor for diffusion through the stomata and boundary layer (ε_*k*_) used in Equation **2** (Mechanism 2). Unsaturated *e*_i_ can change ε_*k*_ in two ways. Firstly, it can increase *g*_s_ (Buckley & Sack 2019), which affects the calculation of ε_*k*_:

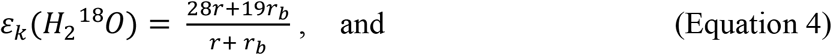

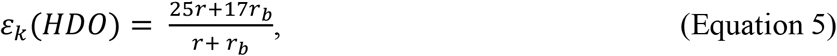

where *r* is stomatal resistance and *r*_b_ is boundary layer resistance (Farquhar *et al*., 1989). A similar calculation for ε_*k*_ has been suggested by Flanagan et al. (1991) (Horita, Rozanski and Cohen, 2008). Since *r* is the inverse of 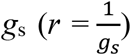 (Horita, Rozanski and Cohen, 2008), an increase in *g*_s_ decreases *r*. Such a decrease in *r* by increased *g*_s_ changes ε_*k*_, and could thus affect LWE.

The second way that unsaturated *e*_i_ can affect ε_*k*_, is based on the understanding that ε_*k*_ represents the nonequilibrium component of leaf water evaporation, where isotope fractionation is controlled by molecular diffusion (Farquhar *et al*., 1989; Flanagan *et al*., 1991; Horita, Rozanski and Cohen, 2008). Such nonequilibrium isotope fractionation has recently been adapted to the specialized marine conditions for evaporation from seawater, for example investigating a turbulent component in response to wind speed (Zannoni *et al*., 2022). Given that ε_*k*_ can be adapted for specialized evaporative conditions, ε_*k*_ has not yet been adapted for unsaturated *e*_i_. Indeed, if *e*_i_ is unsaturated, there would not be ψ equilibrium between apoplastic water and vapor in intercellular spaces, owing to vapor diffusion away from evaporative sites by a small water vapor concentration gradient (Buckley & Sack 2019). Diffusion along a concentration gradient is a source of isotopic fractionation (Merlivat 1978), therefore such diffusion along a concentration gradient within leaf intercellular spaces is a source of isotopic fractionation that could have implications to LWE. Given that Equations **4 & 5** describe kinetic fractionation by stomatal resistance (*r*) and boundary layer resistance (*r*_b_), we suggest incorporating intercellular resistance (*r*_i_) to account for isotope fractionation by diffusion along a vapor concentration gradient within the leaf intercellular space. We suggest that *r*_i_ occurs in Equations **4 & 5** as:

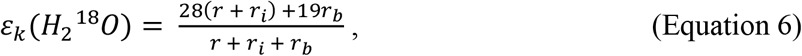

and

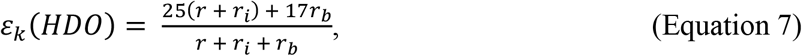

respectively. Here, *r*_i_ is exposed to the same isotope fractionation (28 & 25) as *r*, because they are both characterized by diffusive water vapor molecule movement, and they both contribute to a diffusion layer between an equilibrium layer at the air-water interface, and the boundary layer which is characterized by laminar flow. Since Buckley & Sack (2019) calculated that the water vapor concentration gradient in intercellular spaces would be small, it is unlikely that *r*_i_ is as influential driving factor to LWE compared to, for example, *r*. Nevertheless, since it has not been tested before, it is essential to explore whether an introduction of *r*_*i*_ by unsaturated *e*_i_ is relevant to LWE.

Overall, if *e*_i_ is unsaturated, it can have two effects to LWE. Firstly, it increases the influence of Δ_*v*_ while decreasing the influence of ε_*k*_ (via higher 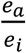 in Equation 2, Mechanism 1). Secondly, it can affect ε_*k*_ by decreasing *r* and introducing *r*_i_ (Equation **6**, **7**, Mechanism 2). Therefore, the main aim of the study was to quantify the effects of unsaturated *e*_i_ to LWE model predictions through testing two hypotheses:

- Hypothesis 1: Unsaturated *e*_i_ increases the influence of Δ_*v*_ and decreases the influence of ε_*k*_ to an extent that is relevant for LWE, shown by a change in LWE predictions in response to a lower RH_cellular_ (Mechanism 1).
- Hypothesis 2: The effect of unsaturated *e*_i_ to LWE by changing ε_*k*_, from decreased *r* and introduced *r*_i_ (Mechanism 2), is not influential to LWE compared to other drivers, such as Mechanism 1.

There are assets to using *in situ* measurements for evaluating the ecological relevance of violated model assumptions, because there are large quantities of data available, and *in situ* measurements provide an ecological perspective to the relative importance of violated model assumptions compared to other sources of error. We firstly tested hypotheses using survey data on Scots pine (*Pinus sylvestris* L.) in a boreal forest. Since LWE changes between species, seasons, and sites, we also applied our analyses to a large-scale dataset from Cernusak et al. (2022) (Snyder *et al*., 2010; Bögelein, Thomas and Kahmen, 2017; Munksgaard *et al*., 2017).

## Materials and Methods

### Field site and sampling

Sampling was conducted at Hyytiälä Forest, which is a managed forest approximately 55 years old, in the southern boreal vegetation zone, southern Finland (61°51’N, 24°17’E, Kolari et al. 2022). It is dominated by Scots pine (*Pinus sylvestris* L.), amongst other species, such as Norway spruce (*Picea abies* (L.) H. Karst), birch (*Betula pendula* Roth, *B. pubescens* Ehrh) and European aspen (*Populus tremula* L.) (Kolari *et al*., 2022). In 2018, the dominant tree height was 23.5m and mean tree height was 19.9m, while tree density was 1304 trees ha^-1^ (Kolari *et al*., 2022). The soil type is Haplic podzol on glacial till, and in most places soil depth is less than 1m, except for moist depressions, which have a thicker layer of soil with a thin layer of peat above them (Kolari *et al*., 2022). Precipitation is distributed somewhat evenly throughout the year and mean annual precipitation between 1981 and 2010 was 711mm (Pirinen *et al*., 2012). Hyytiälä belongs to the Integrated Carbon Observation System (ICOS) network, and a variety of meteorological and leaf gas-exchange parameters are continuously monitored at the site. It is beneficial that there are additional, related tree-physiological investigations from the same site, which can provide deeper insights during data interpretation from this study (Soudant *et al*., 2016; Leppä *et al*., 2022; Tang *et al*., 2022).

Samples were collected between 13:00 and 16:00 during six sampling days with no rain, distributed across the 2019 summer growth season (17 May, 07 June, 28 June, 26 July, 27 August, 23 September). One-year old needles and 2 – 4mm diameter twigs (twig bark was removed) at 18m height were sampled from sun-exposed branches from five Scots pine trees and stored in 12 ml gas-tight glass vials (Exetainer, Labco, UK). All samples were immediately transferred to a cool box. Atmospheric water vapor was collected within the canopy at the same height as needle and twig sampling (18m), on each sampling day, for three hours between 13:00 and 16:00. A dry ice-ethanol cold trap was used, wherein air was pumped into 6mm tubes leading to a U-shaped cold trap (< -70°C) at 0.7 – 1lmin^-1^. The U-tube was then immediately capped tightly, removed, then stored in a cool box. Immediately after fieldwork, the collected moisture was transferred into 2ml IRMS vials using a glass Pasteur pipette and stored in a freezer (-20°C) together with the collected needle and twig samples.

Atmospheric temperature (T_atm_) and RH (RH_atm_) were downloaded from the Smart SMEAR AVAA portal (https://smear.avaa.csc.fi/). They were measured onsite, at the ICOS ecosystem station profile, by a Rotronic MP102H RH/T sensor at 16.8m. Leaf transpiration rate was measured using two automated, box-shaped shoot chamber systems made of acrylic plastic (2.1dm^3^), surrounding debudded shoots in the uppermost canopy (20m, Aalto et al. (2014)). One cuvette monitored one-year old shoots and a second cuvette measured two-year old shoots, and averages from both cuvettes were used. Cuvettes were ventilated and equipped with a fan. Transpiration rate was calculated by applying a non-linear equation to chamber H_2_O vapor concentrations during the first 5 – 35s of intermittent chamber closures (Kolari *et al*., 2012; Leppä *et al*., 2022).

### Laboratory analysis

Water was cryogenically extracted from needles and twigs at the Swiss Federal Institute for Forest, Snow and Landscape Research (WSL) (West, Patrickson and Ehleringer, 2006).

Stable isotope analyses were conducted at The University of Basel Stable Isotope Ecology Laboratory, Switzerland, by Thermal Conversion / Elemental Analyzer (TC/EA) coupled to a Delta V Plus isotope ratio mass spectrometer (IRMS) through a ConFlo IV interface (Thermo Fisher Scientific, Bremen, Germany) (Newberry, Nelson and Kahmen, 2017). Samples were injected at least six times, and a minimum of three of the measurements were used to calculate a mean value, since starting measurements were omitted to compensate for memory effects from the previous sample. Measurements were normalized to Vienna Standard Mean Ocean Water (VSMOW) using calibrated in-house standards with a δ^2^H value of -76.4‰, and a δ^18^O value of -10.7‰. Isotope values were defined as:

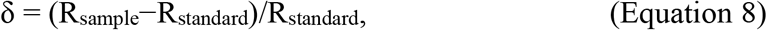

relative to VSMOW, where R is the D/H or 18O/16O ratio for δ^2^H and δ^18^O, respectively. The standard deviations of quality controls during the time of analyses were 0.3‰ (n = 49) for δ^2^H and 0.12‰ (n = 49) for δ^18^O.

### Large-scale dataset sourcing

The large-scale dataset and its LWE predictions were first sourced from the review published by Cernusak et al. (2022). This large-scale dataset comprises of 546 datapoints for paired Δ^2^H_lw_ and Δ^18^O_lw_. The geographical range extends across more than 100º of latitude and there is an elevation range larger than 3000m. Most of the data is from temperate forests or woodlands, followed by tropical forests or woodlands. The data from Hyytiälä was added to the large-scale dataset and, after a grassland in Greenland, it contributed the highest-latitude data and the only boreal forest measurements. The large-scale dataset was filtered to select sampling sites with at least five different sampling times, to meet statistical analysis criteria. Data from Kahmen et al. (2011) were clustered into five main sampling sites, and three sites from Munksgaard et al. (2017) were clustered into one site (Herberton, Wild River & Mount Garnet), so that data met the filter criterion and thus could be included. The resultant dataset constituted of 534 datapoints from Cernusak et al. (2022) and 29 datapoints from this Hyytiälä (∑ = 563).

### Leaf water heavy isotope enrichment modelling

All modelling and statistical analyses were performed in R (R Core Team, 2022). Observed leaf water and water vapor enrichments were calculated as:

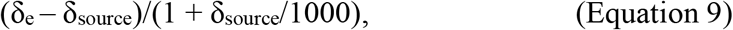

where δ_e_ is the isotope value of the parameter whose enrichment above source water is being estimated, i.e., leaf water or water vapor (Cernusak *et al*., 2016). Firstly, LWE was modelled using Equation **2**, using the calculations provided in the supporting materials by Cernusak et al. (2022), for both Hyytiälä and the large-scale dataset. At Hyytiälä, T_leaf_ was first assumed to be the same as T_atm_, which is a reasonable assumption because the Scots pine needles are small and well-coupled to the atmosphere (Launiainen *et al*., 2016; Kim *et al*., 2018; Leppä *et al*., 2022). Nevertheless, given that there are uncertainties relating to the assumption that T_leaf_ is equal to T_atm_, and that recent evidence shows that the relationship between T_leaf_ and T_atm_ can change on a diurnal basis (Still *et al*., 2022), we expanded analyses to include sensitivity of results to a T_leaf_ change of ±2ºC from T_atm_, to guide inferences on the relative influence of unsaturated *e*_i_ to LWE compared to a ±2ºC change in T_leaf_.

Main results were inferred from the foundational CG model (Equation **2**). A two-pool correction and a Péclet correction were additionally applied, as an initial demonstration of how such corrections can interact with unsaturated *e*_i_. The two-pool correction was calculated as:

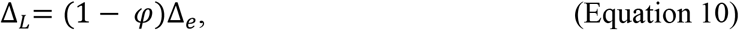

where Δ_*L*_ is the final calculated leaf water heavy isotope enrichment (LWE), Δ_*e*_ is the modelled LWE by the Craig-Gordon model, and φ is the proportion of unenriched xylem water in leaf water (Leaney *et al*., 1985; Song *et al*., 2015). The estimate for φ was 0.316, based on Scots pine leaf anatomical measurements by Roden et al. (2015). For the Péclet correction, the Péclet number was calculated as:

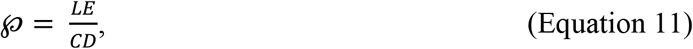

where *L* is effective path length, *E* is transpiration rate (mol m^-2^ s^-1^), *C* is the molar concentration of water (5.5 ×10^-4^ mol m^-3^) and *D* is the diffusivity of the water isotopologue responsible for enrichment. Their calculation is described in further detail by Cernusak et al. (2016), and in this study, *L* was calculated for each sampling date based on transpiration rate. The Péclet correction is applied as *P*:

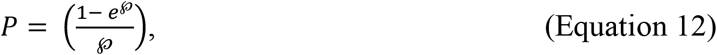

where:

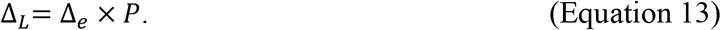

There was limited transpiration rate and φ data availability for the large-scale dataset, so this additional model correction demonstration was only performed for Hyytiälä data.

To explore the effects of unsaturated *e*_*i*_ to LWE by increasing influence of Δ_*v*_ and decreasing influence of ε_k_ (Hypothesis 1), Equation **2** was applied with different assumptions for RH_cellular_ when calculating *e*_i_. For example, in Equation **2**, atmospheric vapor pressure (*e*_a_) can be expressed as:

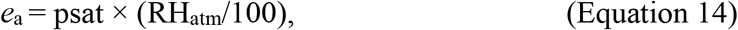

where psat is saturated vapor pressure. Since *e*_i_ was assumed to be saturated during LWE modelling, it has been estimated as *e*_i_ = psat. In this study, we calculated *e*_i_ in the same way that *e*_a_ has been expressed, in Equation **14**, by replacing RH_atm_ with RH_cellular_:

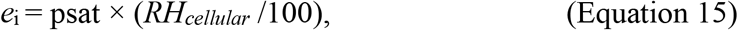

using the following assumptions for RH_cellular_:

1. RH_cellular_ = 100%. *Saturated e*_i_.
2. RH_cellular_ = 90%. *Within the observed range reported by literature (Cernusak et al., 2018; Wong et al., 2022)*.
3. RH_cellular_ = 80%. *The lowest approximate RH*_*cellular*_ *reported by literature (Cernusak et al., 2018; Wong et al., 2022)*.

Finally, since RH_atm_ potentially affects RH_cellular_ (Vesala *et al*., 2017; Cernusak *et al*., 2018), we modelled RH_cellular_ as a response to RH_atm_. This was a model-optimization, which used measured LWE to find a fitted *e*_i_ along an RH_atm_ gradient for both Δ^2^H_lw_ and Δ^18^O_lw_. We used:

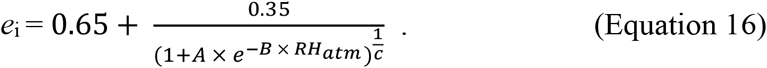

Calculants A, B and C were solved simultaneously for both Δ^2^H_lw_ and Δ^18^O_lw_, to find one fitted RH_cellular_ for both elements using the optim function in the ‘stats’ package. The optim function was run with default configuration using the Nelder-Mead algorithm. Fitted RH_cellular_ was also solved for each of Δ^2^H_lw_ and Δ^18^O_lw_ separately.

When calculating psat for *e*_a_, T_atm_ is used, meanwhile, when calculating psat for *e*_i_, T_leaf_ is used (Cernusak *et al*., 2016). In Equation **2**, *e*_i_ occurs twice, and main results are given for adjustment of both *e*_i_ occurrences. An additional post-hoc analysis was performed on data from Hyytiälä, where each occurrence of *e*_i_ was adjusted to different RH_cellular_ assumptions, separately.

To test the effect of unsaturated *e*_i_ to isotope fractionation associated with *r*_i_, we calculated ε_k_ using Equations **6 & 7** and implemented the altered ε_k_ to Equation **2**, to look for observable changes in predicted LWE compared to modelled LWE using ε_k_ that had been calculated using Equations **4 & 5**. For this, we used data from Hyytiälä when RH_cellular_ was 90% or 80%, and we applied leaf anatomical measurements of Scots Pine needles by Roden et al. (2015). Intercellular resistance (*r*_i_) was estimated using three calculation steps. First, the rate of water vapor diffusion along a concentration gradient within an intercellular space (*J*, mol m^-2^ s^-1^) was estimated using Fick’s law of diffusion:

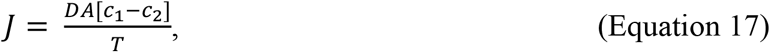

where *D* is the diffusion constant, at 2.44×10^-5^ m^-2^ s^-1^ (Merlivat 1978), *A* is the cross-sectional area of diffusion, which we approximated by using cross-sectional leaf area exposed to evaporation per square meter using Scots Pine measurements by Roden et al. (2015) (*A* = 1 -φ = 0.684). Then, *c*_1_ was the concentration of water vapor in the equilibrium layer at the air-water interface in the leaf intercellular space (mol m^-3^), calculated as saturated vapor concentration at leaf temperature, and *c*_2_ was the unsaturated concentration of water vapor in the unsaturated portion of the leaf intercellular space (*c*_2_ = *c*_1_ × (RH_cellular_/100)). Such definitions of *c*_1_ and *c*_2_ were based on the principle that there is an equilibrium layer at the air-water interface during evaporation, and because there are humidity gradients inside of leaf intercellular spaces (Wong et al. 2022), but they can be improved if more knowledge arises about leaf intercellular space humidity conditions. Finally, *T* was the length of the diffusion pathway, which has not yet been quantified, we approximated *T* by using measured mean mesophyll thickness for Scots Pine by Roden et al. (2015) (1.71×10^-4^ m).

Intercellular conductance (*g*_i_, mol m^-2^ s^-1^) was then calculated using an adaptation of the following calculation:

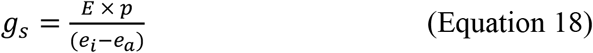

where g_s_ is stomatal conductance (mol m^-2^ s^-1^), *E* is transpiration rate (mol m^-2^ s^-1^), p is air pressure (kPa), *e*_a_ is atmospheric water vapor pressure (kPa) and *e*_i_ is water vapor pressure in the leaf intercellular space (kPa). We adapted the equation to:

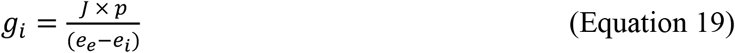

where *e*_e_ is water vapor pressure at the equilibrium layer of the air-water interface in the leaf intercellular space (kPa), calculated as saturated vapor pressure at leaf temperature. Then, *e*_i_ (kPa) was adjusted to the tested level of unsaturation within the leaf intercellular space (*e*_i_ = *e*_e_ × (RH_cellular_/100)). We then calculated intercellular resistance (mol m^-2^ s^-1^) as:

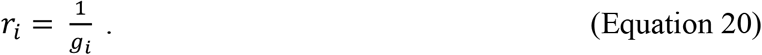

During the calculations, we assumed that the length of the diffusion pathway was equal to mean mesophyll thickness, and that the cross-sectional area of diffusion was equal to the cross-sectional area of a leaf exposed to evaporation, therefore *r*_i_ estimates were approximate. However, given that the *r*_i_ response to unsaturated *e*_i_ varies along a much smaller magnitude than the variability of *r*, the consequences of the described assumptions are unlikely consequential to this study, where *r*_i_ has been added to *r* when calculating ε_k_ (Equation **6**, **7**).

### Statistical analyses

At Hyytiälä and in the large-scale dataset, linear mixed models (LMMs) with random intercepts were used to compare modelled to observed LWE. At Hyytiälä, the random intercept was sampling date, while in the larger dataset the random intercept was site ID with rank sampling time nested inside of site ID. Unadjusted Intraclass Correlations (ICC) were used to quantify unexplained variability between random factors which remained after modelled LWE was compared to observed LWE (Nakagawa, Johnson and Schielzeth, 2017). One outlier leaf water δ^18^O measurement was removed from Hyytiälä data. Each LMM analysis was accompanied with a calculation of Root Mean Square Error (RMSE), which is an estimate of overall proximity of predicted LWE to observed LWE.

## Results

### Data Overview

Data used as input to the Craig-Gordon model to predict LWE at Hyytiälä were within the (Still *et al*., 2022) range of the large-scale dataset (Fig. **1a-d**). Their means were lower at Hyytiälä compared to the large-scale dataset, most noticeably so for ^2^H enrichment of water vapor above source water, which was 11.4‰ lower at Hyytiälä than in the large-scale dataset (Δ^2^H_wv_, Fig. **1c**). Congruently, observed LWE at Hyytiälä was within the range of observed LWE in the large-scale dataset (Fig. **1e-f**). Nevertheless, mean observed Δ^2^H_lw_ was approximately the same at Hyytiälä and in the large-scale dataset, while mean observed Δ^18^O_lw_ was 2.4‰ higher at Hyytiälä than in the large-scale dataset (Fig. **1e-f**). The seasonal variability of leaf water isotope enrichment at Hyytiälä covered a substantial proportion of the data range in the large-scale dataset, at 34% for each of Δ^2^H_lw_ and Δ^18^O_lw_.

**Figure 1.**
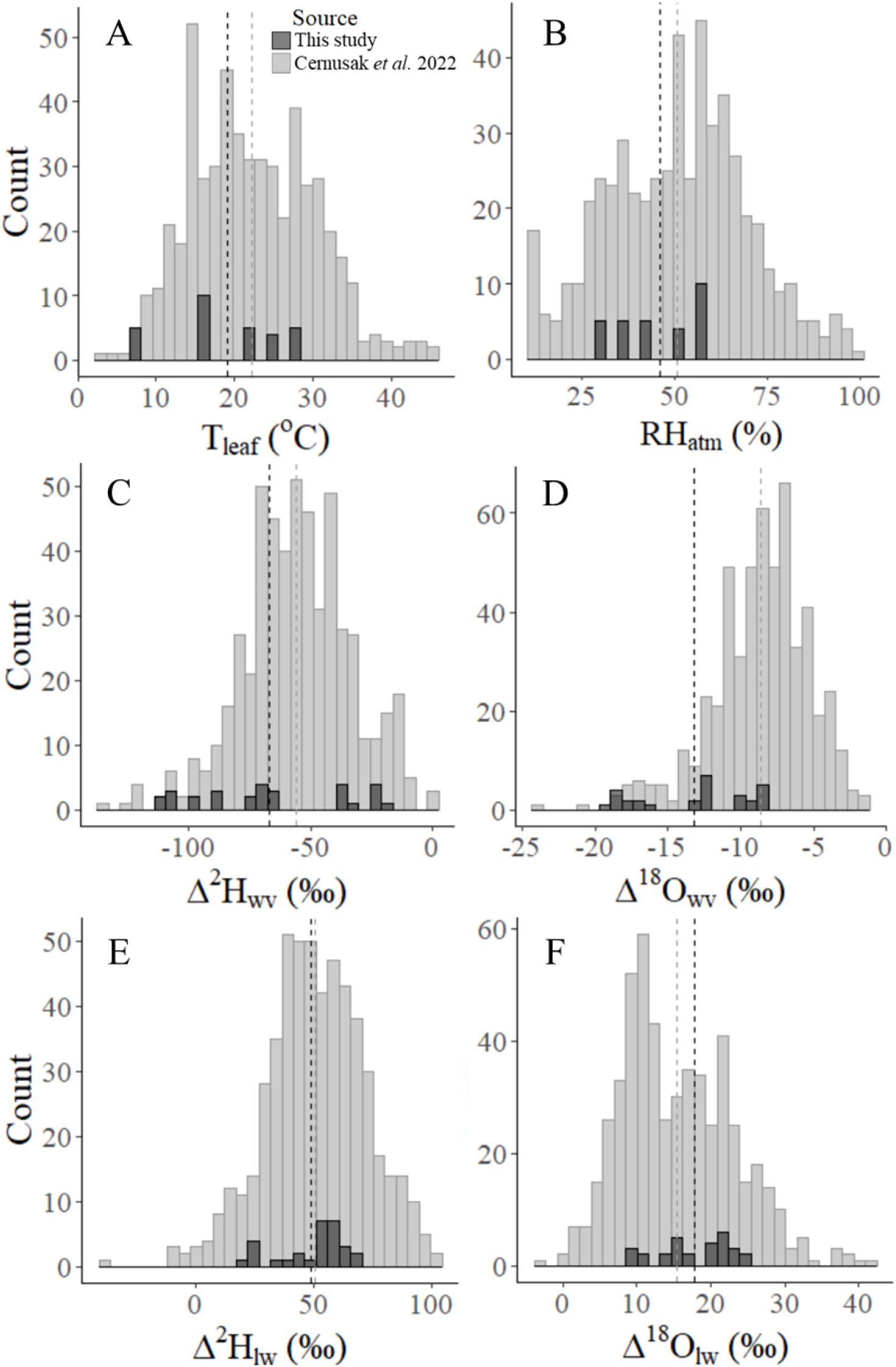
Frequency distributions of leaf temperature (T_leaf_ (ºC), A), atmospheric relative humidity (RH_atm_ (%), B), water vapor deuterium (Δ^2^H_wv_ (‰), C) and oxygen-18 (Δ^18^O_wv_ (‰), D) enrichment above source water, and observed leaf water deuterium (Δ^2^H_lw_ (‰), E) and oxygen-18 (Δ^18^O_lw_ (‰), F) enrichment above source water, n = 563. Data from this study (dark grey) shows seasonal variability for *P. sylvestris* during the 2019 growing season at Hyytiälä, Finland, and it overlays a selection of review data collected by Cernusak et al. (2022, lighter grey). Dashed lines show the mean of each parameter, for Hyytiälä and the review data.

### Hypothesis 1: Increased influence of Δ_*v*_ and decreased influence of ε_k_, by unsaturated *e*_i_, is relevant to LWE

#### Hyytiälä

##### Foundational Craig-Gordon model

When assumed RH_cellular_ was lowered from 100% to 90% and 80%, predicted Δ^2^H_lw_ and Δ^18^O_lw_ became noticeably lower (red series in Fig. **2a-c**; Table **2**). This improved Δ^2^H_lw_ predictions by reducing the offset between observed and modelled values, because the predictions based on 100% RH_cellular_ largely overestimated Δ^2^H_lw_ (red series in Fig. **2a**; Table **2**). However, for Δ^18^O_lw_, 100% RH_cellular_ already provided a good agreement between the measured and modelled Δ^18^O_lw_, with only a modest average model overestimation of 1‰ (red series in Fig. **2e**). Hence, the lowering of RH_cellular_ to 90% or 80% led to increased error of Δ^18^O_lw_ predictions, by underestimation (red series in Fig. **2f-g**; Table **2**). The accuracy of LWE predictions was affected by ±2ºC variability in T_leaf_ more for Δ^18^O_lw_ than for Δ^2^H_lw_, and the impact was larger on lower enrichments, for predictions by models with 100%, 90% and 80% RH_cellular_ (horizontal lines in Fig. **2a-c, e-g**).

**Figure 2.**
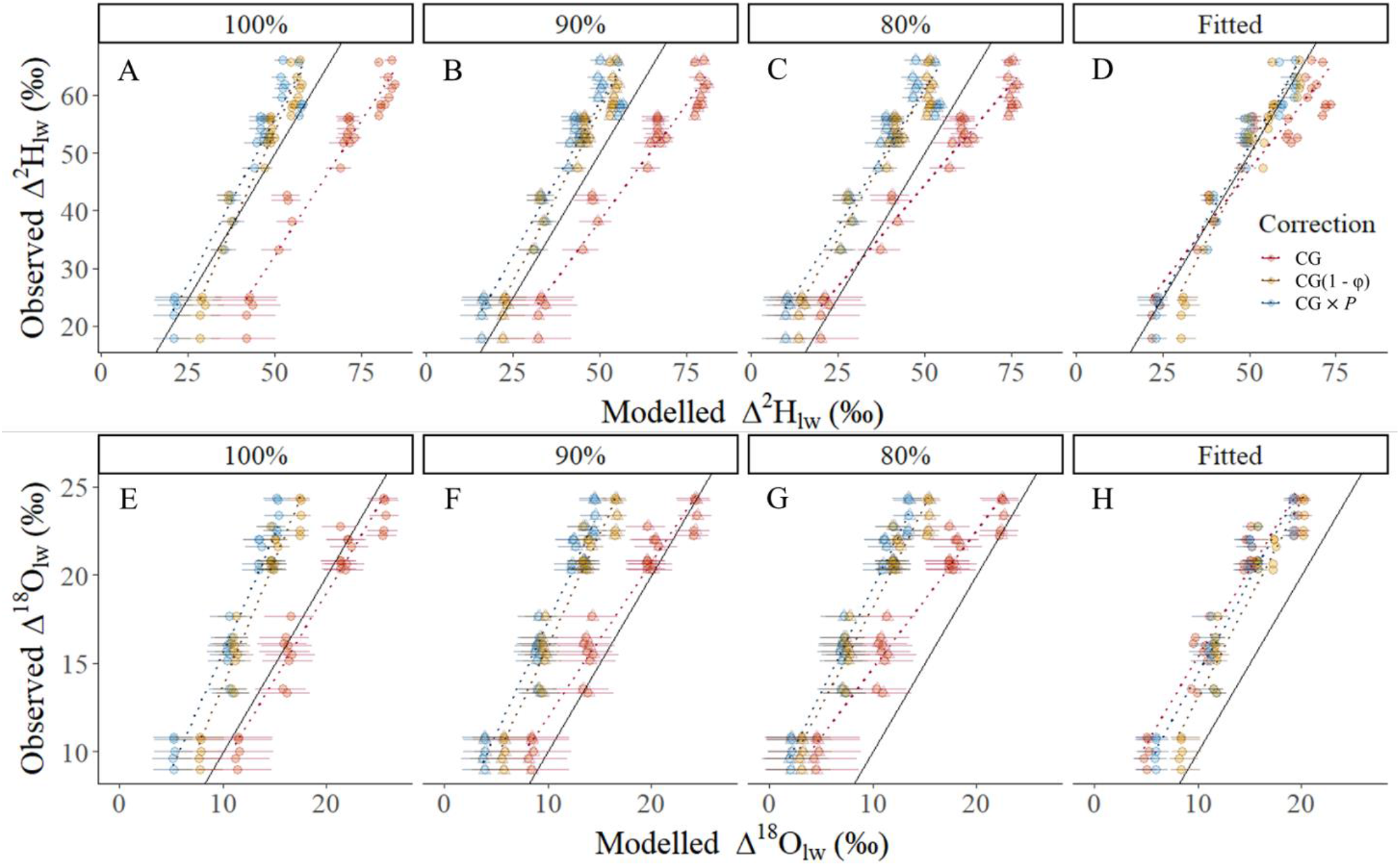
Relationships between modelled and measured Hyytiälä leaf water deuterium (Δ^2^H_lw_) and oxygen-18 (Δ^18^O_lw_) enrichment, when leaf intercellular space relative humidity (RH_cellular_) was changed in the models (100%, 90%, 80%, fitted RH_cellular_. n = 29), to test unsaturated *e*_i_ effects to Δ^2^H_lw_ and Δ^18^O_lw_ via increased influence of Δ_*v*_ and decreased influence of ε_k_ to LWE. Dashed lines show linear mixed model fits, solid black lines demonstrate a 1:1 relationship, and horizontal lines show model variability in response to ±2ºC leaf temperature. Triangles in graphs with 90% and 80% RH_cellular_ show model results once intercellular resistance (*r*_i_) has been included in the calculation of ε_k_. CG: Craig-Gordon model; CG(1 – φ): Craig-Gordon model with two-pool correction; CG × *P*: Craig-Gordon model with Péclet correction.

When RH_cellular_ was reduced to 90% or 80%, the lower predicted enrichments were lowered to a larger extent than higher predicted enrichments, as indicated by the increase in intercepts and the decline in slopes, for both elements (Fig. **2a-c, e-g**; Table **2**). This attribute meant that, while reductions in RH_cellular_ could reduce model prediction offsets from observed values if a model otherwise overestimated LWE (Δ^2^H_lw_), they had a biased influence on model prediction accuracy. Such a prediction accuracy bias was completely remediated for Δ^18^O_lw_, by the model optimization that found a fitted RH_cellular_ that varied along a RH_atm_ gradient, albeit with an offset between observed and measured values (Fig. **2 h**; Table **2**). Meanwhile, for Δ^2^H_lw_ predictions, the prediction accuracy bias was only partly remediated by fitted RH_cellular_, because it improved the prediction accuracy bias compared to 80% RH_cellular_, but it worsened the prediction accuracy bias compared to 90% RH_cellular_ (Table **2**).

The fitted-RH_cellular_ for Δ^18^O_lw_ consistently underestimated observed Δ^18^O_lw_, for the foundational CG model (Fig. **2 h**, Table **2**). Indeed, the Craig Gordon model assuming 100% RH_cellular_ remained the best predictor of Δ^18^O_lw_ (Fig. **2 e**, Table **2**). In contrast, for Δ^2^H_lw_, the fitted RH_cellular_ exhibited reduced offsets between modelled and measured values, producing better Δ^2^H_lw_ predictions compared to non-fitted RH_cellular_, observed by a lowered Root Mean Square Error (RMSE) (Fig. **2 d**; Table **2**).

Performance of LWE models deteriorated extremely when only one of the two *e*_i_ occurrences in Equation **2** was adjusted for unsaturated *e*_i_ (Supplemental Table **1**), showing that the relationships between *e*_i_ and both Δ_*v*_ and ε_k_, are important to the response of LWE to unsaturated *e*_i_.

##### Péclet and two-pool corrections

Main results from this study can be derived from the foundational CG model, and results of additional Péclet and two-pool corrections are described to demonstrate how such corrections might interact with a decrease in *e*_i_ in Equation **2** tested for Hypothesis 1. Therefore, we used one calculation of effective path length for the Péclet correction, and a literature-derived constant φ for the two-pool correction.

The two-pool and Péclet correction almost always lowered Δ^2^H_lw_ and Δ^18^O_lw_ predictions compared to the CG model and they had larger effects at higher enrichments, except for when RH_cellular_ was fitted (orange series in Fig. **2**; Table **2**). Resultantly, they mostly underestimated Δ^2^H_lw_ and Δ^18^O_lw,_ but they still improved Δ^2^H_lw_ predictions when RH_cellular_ was 100%, 90%, or fitted (RMSE in Table 2). The Péclet correction had less of an effect bias to higher enrichments compared to the two-pool correction (blue series in Fig. **2**; Table **2**). Since the two-pool correction, the Péclet correction and the lowered RH_cellular_ all typically lowered predicted Δ^2^H_lw_ and Δ^18^O_lw_, when the two-pool or Péclet correction were combined with lowered RH_cellular_, they led to even lower predicted Δ^2^H_lw_ and Δ^18^O_lw_ than if applied individually, except for when RH_cellular_ was fitted (Fig. **2**; Table **2**). Indeed, Δ^2^H_lw_ was almost perfectly predicted, when RH_cellular_ was fitted after a Péclet correction had been applied (Fig. **2 d**; Table **2**). However, unrealistically high fitted RH_cellular_ were needed for the model optimization (103 – 146% and 105 – 210%, respectively; Supplemental Fig. **1**), showing that the versions of two-pool and Péclet corrections used in this study were thus fundamentally incompatible with reductions in RH_cellular_ at Hyytiälä. Nevertheless, the larger effect to higher enrichments by the two-pool and Péclet corrections balanced the larger effect to lower enrichments by RH_cellular_ at 90% or 80%, thus reducing resultant model prediction accuracy bias at different LWE, despite frequent underestimations.

**Table 2.** Linear mixed model fits between modelled and observed leaf water deuterium (Δ^2^H_lw_) and oxygen-18 (Δ^18^O_lw_) enrichment at Hyytiälä (n = 29), with models using different intercellular relative humidity (RH_cellular_, %, Mechanism 1. CG: Craig-Gordon model; CG(1 - φ): Craig-Gordon model with two-pool correction; CG × *P*: Craig-Gordon model with Péclet correction; R^2^(M): Marginal R^2^; R^2^(C): Conditional R^2^; ICC: Intraclass correlation between sampling dates).

| | Model correction | $\text{RH}_{\text{cellular}}$ (%) | Intercept | Slope<sup>a</sup> | ICC | $R^2(\text{M})$ | $R^2(\text{C})$ | RMSE |
| --- | --- | --- | --- | --- | --- | --- | --- | --- |
| Hyytiälä<br>$\Delta^2\text{H}_{\text{lw}}$ | CG | 100 | $-14.26 \pm 3.88$ | **$0.93 \pm 0.06$** | 0.01 | 0.95 | 0.96 | 19.27 |
| | CG(1 - $\varphi$) | | $-14.26 \pm 3.88$ | **$1.36 \pm 0.08$** | 0.01 | 0.95 | 0.96 | 5.44 |
| | CG × $P$ | | $-1.29 \pm 4.55$ | **$1.14 \pm 0.1$** | 0.03 | 0.93 | 0.96 | 6.37 |
| | CG | 90 | $-2.26 \pm 2.67$ | **$0.81 \pm 0.04$** | 0.01 | 0.95 | 0.96 | 14.52 |
| | CG(1 - $\varphi$) | | $-2.26 \pm 2.67$ | **$1.19 \pm 0.06$** | 0.01 | 0.95 | 0.96 | 7 |
| | CG × $P$ | | $5.91 \pm 4$ | **$1.05 \pm 0.09$** | 0.03 | 0.92 | 0.96 | 8.87 |
| | CG | 80 | $9.29 \pm 2.01$ | **$0.7 \pm 0.03$** | < 0.005 | 0.95 | 0.96 | 9.94 |
| | CG(1 - $\varphi$) | | $9.29 \pm 2.01$ | **$1.03 \pm 0.05$** | < 0.005 | 0.95 | 0.96 | 10.7 |
| | CG × $P$ | | $13.48 \pm 3.44$ | **$0.95 \pm 0.09$** | 0.04 | 0.92 | 0.96 | 12.51 |
| | CG | Fitted | $8.46 \pm 4.67$ | **$0.77 \pm 0.09$** | 0.06 | 0.9 | 0.96 | 7.24 |
| | CG(1 - $\varphi$) | | $-9.79 \pm 6.06$ | **$1.18 \pm 0.12$** | 0.05 | 0.91 | 0.96 | 4.79 |
| | CG × $P$ | | $-0.45 \pm 3.08$ | **$1.03 \pm 0.06$** | 0.01 | 0.95 | 0.96 | 3.48 |
| Hyytiälä<br>$\Delta^{18}\text{O}_{\text{lw}}$ | CG | 100 | $-0.58 \pm 1.12$ | **$0.98 \pm 0.06$** | 0.01 | 0.95 | 0.96 | 1.51 |
| | CG(1 - $\varphi$) | | $-0.58 \pm 1.12$ | **$1.43 \pm 0.08$** | 0.01 | 0.95 | 0.96 | 5.26 |
| | CG × $P$ | | $2.32 \pm 1.4$ | **$1.33 \pm 0.12$** | 0.03 | 0.93 | 0.96 | 6.4 |
| | CG | 90 | $3.18 \pm 0.86$ | **$0.88 \pm 0.05$** | 0.01 | 0.95 | 0.96 | 1.68 |
| | CG(1 - $\varphi$) | | $3.18 \pm 0.86$ | **$1.28 \pm 0.07$** | 0.01 | 0.95 | 0.96 | 6.57 |
| | CG × $P$ | | $4.61 \pm 0.94$ | **$1.27 \pm 0.09$** | 0.02 | 0.94 | 0.96 | 7.59 |
| | CG | 80 | $6.91 \pm 0.63$ | **$0.78 \pm 0.04$** | 0.01 | 0.95 | 0.96 | 4.12 |
| | CG(1 - $\varphi$) | | $6.91 \pm 0.63$ | **$1.13 \pm 0.06$** | 0.01 | 0.95 | 0.96 | 8.3 |
| | CG × $P$ | | $7.28 \pm 0.57$ | **$1.2 \pm 0.06$** | < 0.005 | 0.95 | 0.96 | 9.12 |
| | CG | Fitted | $5.46 \pm 1.02$ | **$0.99 \pm 0.08$** | 0.02 | 0.93 | 0.96 | 5.44 |
| | CG(1 - $\varphi$) | | $1.74 \pm 1.56$ | **$1.13 \pm 0.11$** | 0.04 | 0.92 | 0.96 | 3.9 |
| | CG × $P$ | | $3.68 \pm 1.24$ | **$1.07 \pm 0.09$** | 0.03 | 0.93 | 0.96 | 4.88 |

<sup>a</sup>Significant relationship when slope is **bold** ( $p < .001$ ).

#### Large-scale dataset

Results from the large-scale dataset were mostly parallel to results from Hyytiälä. Like at Hyytiälä, a reduction in RH_cellular_ from 100%, to 90% and 80%, led to less enriched Δ^2^H_lw_ and Δ^18^O_lw_ predictions (Fig. **3**; Table **3**). Such reductions in predicted Δ^2^H_lw_ and Δ^18^O_lw_ clearly benefited Δ^2^H_lw_ prediction accuracy by reducing overestimates, most evidently shown by a large decrease in RMSE of model predictions (Table **3**). These outcomes are the same as results observed for Hyytiälä (Table **2**). However, Δ^18^O_lw_ prediction accuracy improved when RH_cellular_ was lower than 100%, as shown by a decrease in RMSE (Table **3**), whereas Δ^18^O_lw_ prediction accuracy decreased when RH_cellular_ was lower than 100% at Hyytiälä (Table **2**). The larger decrease in predicted LWE at lower predicted enrichments when RH_cellular_ was lowered from 100% to 90% or 80%, persisted beyond the Hyytiälä dataset to the large-scale dataset, for both elements (Fig. **3**; Table **3**). Like for Hyytiälä, this bias was mostly remediated for Δ^18^O_lw_ by using a fitted RH_cellular_ based on observed LWE and RH_atm_ (Fig. **3**; Table **3**). However, the fitted RH_cellular_ also largely remediated the bias for Δ^2^H_lw_, unlike at Hyytiälä (Fig. **3**; Table **3**). In congruence with findings from Hyytiälä, the best-fitting model for Δ^2^H_lw_ was the model with a fitted RH_cellular_, while unlike at Hyytiälä, Δ^18^O_lw_ was best explained by a model assuming 90% RH_cellular_ rather than 100% RH_cellular_. Overall, results from the large-scale dataset reinforce the observed relevance of reduced RH_cellular_ to LWE observed at Hyytiälä, because they demonstrate that predictions of both Δ^2^H_lw_ and Δ^18^O_lw_ noticeably change, and even improve, in response to unsaturated *e*_i_.

**Figure 3.**
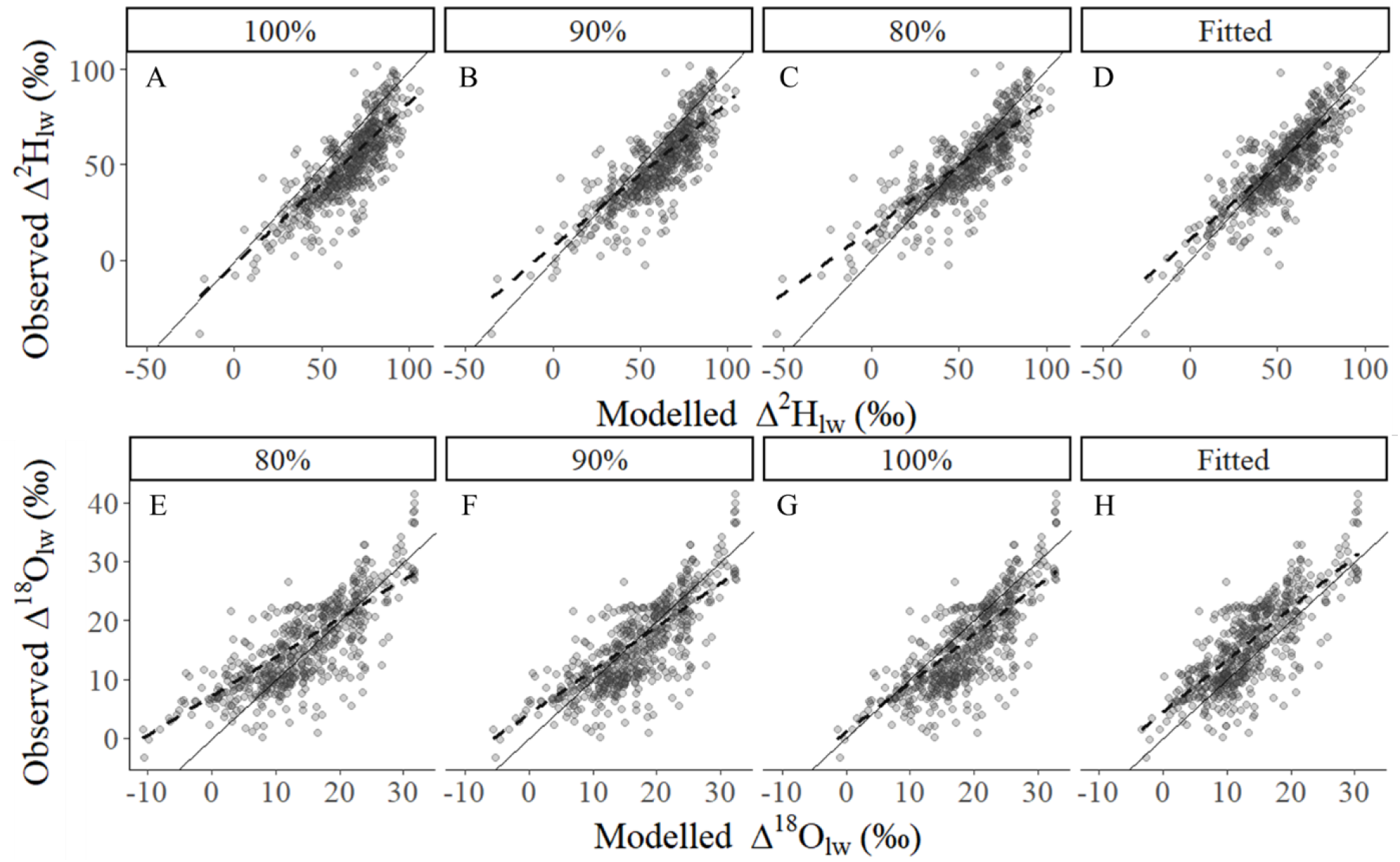
Relationships between Craig-Gordon model predictions and measured leaf water deuterium (Δ^2^H_lw_) and oxygen-18 (Δ^18^O_lw_) enrichment with different assumptions of leaf intercellular space relative humidity (100%, 90%, 80%, fitted), in the studied large-scale dataset. The dataset includes data from this study combined with review data from Cernusak et al. (2022) (Mechanism 1, n = 563). Dashed lines show linear mixed model fits and solid lines demonstrate the 1:1 relationship.

**Table 3.**
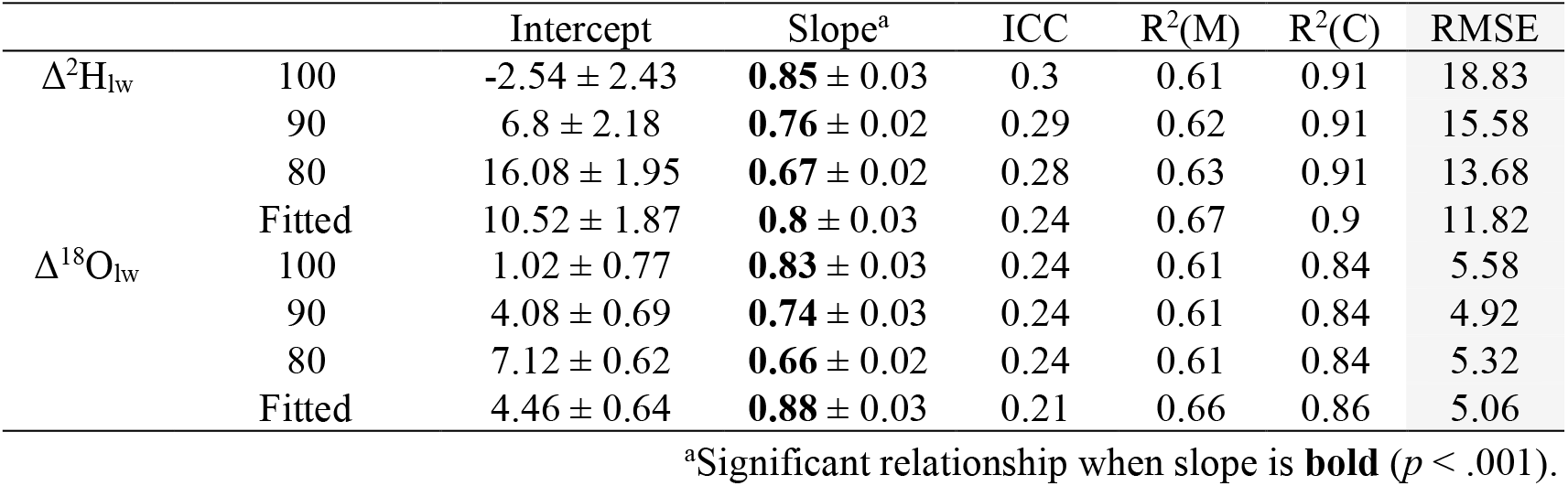
Linear mixed model fits between modelled and observed leaf water deuterium (Δ^2^H_lw_) and oxygen-18 (Δ^18^O_lw_) enrichment in the studied large-scale dataset, with models using different intercellular relative humidity (RH_cellular_ (%), Mechanism 1, n = 563).

#### Fitted RH_cellular_ predictions

The fitted RH_cellular_ increased as RH_atm_ increased (Fig. **4**). The fitted RH_cellular_ for *P. sylvestris* at Hyytiälä was highly complementary to the fitted RH_cellular_ for the larger dataset, as indicated by the almost overlapping fitted RH_cellular_ along the common RH_atm_ gradient. Sensitivity tests for ±2ºC change in T_leaf_ showed that fitted RH_cellular_ is influenced by ±2ºC changes in T_leaf_, at both Hyytiälä and in the larger dataset (shaded regions in Fig. **4**).

**Figure 4.**
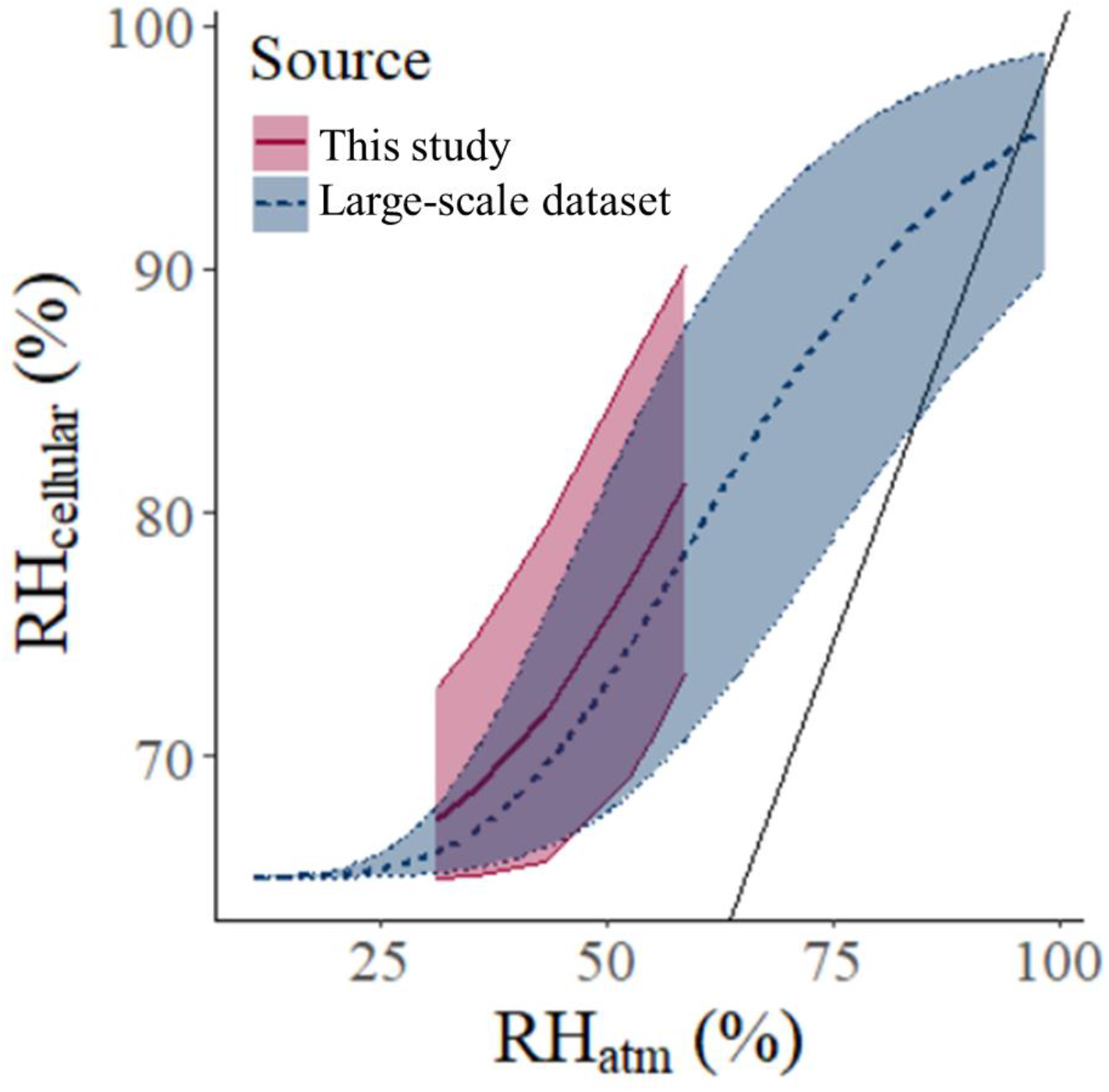
Fitted intercellular relative humidity (RH_cellular_) in response to atmospheric RH (RH_atm_) for *Pinus sylvestris* at Hyytiälä, Finland (n = 29, “This study”), and for the large-scale dataset which combines data from this study with review data from Cernusak et al. (2022) (n = 563, “Large-scale dataset”). The solid line demonstrates the 1:1 relationship, and shaded areas show fitted RH_cellular_ sensitivity to ±2°C leaf temperature.

### Hypothesis 2: Intercellular resistance fractionation irrelevant to LWE

At Hyytiälä, the isotope fractionation by decreased *r* and introduced intercellular resistance (*r*_i_) from unsaturated *e*_i_ had a negligible influence on predicted Δ^2^H_lw_ and Δ^18^O_lw_, at RH_cellular_ 90% and 80% (triangles strongly overlapped by circles in Figure 3 B, C, F & G). It changed ε_k_ estimates by less than 0.26 and 0.29 for Δ^2^H_lw_ and Δ^18^O_lw_, respectively, for 90% and 80% RH_cellular_. Resultantly, it changed LWE by only 0 – 0.17‰ at 90% and 80% RH_cellular_ for both Δ^2^H_lw_ and Δ^18^O_lw_.

## Discussion

This study is the first to quantitatively evaluate the ecological relevance of unsaturated *e*_i_ to LWE. Overall, results showed that unsaturated *e*_i_ effects is likely relevant to LWE by changing LWE predictions, via both increased influence of Δ_*v*_ and decreased influence of ε_k_ (Fig. **2**, **3**; Table **2**, **3**, Supplemental Table **1**). This means that it is necessary to consider unsaturated *e*_i_ as an important source of error to LWE predictions and reconstructions from organic material, albeit one which can be corrected. In this study, such corrections to *e*_i_ clearly benefited Δ^2^H_lw_ predictions, and conditionally benefited Δ^18^O_lw_ predictions (Fig. **2**, **3**; Table **2**, **3**). Results suggested that additional fractionation by concentration-driven diffusion in leaf intercellular spaces (Equation **6**, **7**) is unlikely relevant to LWE.

### Correction for unsaturated *e*_i_ in studies that use leaf water isotopes

Overall, when RH_cellular_ and thus *e*_i_ was lowered in the foundational CG model, both Δ^2^H_lw_ and Δ^18^O_lw_ predictions changed (Fig. **2**, **3**; Table **2**, **3**). Such changes improved Δ^2^H_lw_ predictions produced by the CG model because the offset between observed and measured Δ^2^H_lw_ decreased, at both Hyytiälä and in the large-scale dataset (RMSE in Table **2**, **3**). Meanwhile, the benefits of lowered RH_cellular_ were not clear for Δ^18^O_lw_ because all reductions in RH_cellular_ in the CG model had effects too large for Δ^18^O_lw_ predictions at Hyytiälä, while all reductions in RH_cellular_ in the CG model improved Δ^18^O_lw_ predictions in the large-scale dataset compared to the CG model assuming saturated *e*_i_ (RMSE in Table **2**, **3**). The more evident benefit to Δ^2^H_lw_ could have been because Δ^2^H_lw_ can be strongly related to the isotopic disequilibrium between water vapor and source water, which was changed by the reductions in RH_cellular_ tested in this study, while Δ^18^O_lw_ is more strongly related to RH_atm_ (Munksgaard *et al*., 2017; Cernusak *et al*., 2022). These results show how it is potentially valuable to account for unsaturated *e*_i_ during Δ^2^H_lw_ predictions and Δ^2^H_lw_ reconstructions from plant compounds, such as tree rings or *n*-alkanes, because unsaturated *e*_i_ directly affects the factors known to be most strongly related to Δ^2^H_lw_.

When using a constant lowered RH_cellular_ (90% or 80%), model bias increased, because the reduced RH_cellular_ affected lower predicted LWE more than higher predicted LWE for both Δ^2^H_lw_ and Δ^18^O_lw_ (Fig. **2**, **3**; Table **2**, **3**). It is noteworthy to recognize that this means that when 100% RH_cellular_ is being used when intercellular spaces are unsaturated, then it thus brings a model prediction accuracy bias of its own. This shows that it is valuable to start using a variable RH_cellular_ along a range of LWE values when calculating *e*_i_, which is supported by evidence that RH_cellular_ changes (Cernusak *et al*., 2018; Wong *et al*., 2022). When results from this study are combined with measurements of RH_cellular_ responses to VPD from Cernusak et al. (2018), a viable solution for estimating RH_cellular_ is calculating RH_cellular_ from RH_atm_ or atmospheric VPD, which are negatively correlated to one another. More studies like Cernusak et al. (2018) are required to gather species-specific RH_cellular_ responses to changing RH_atm_ or VPD, their study can be used to tentatively estimate *e*_i_ of two species: *Juniperus monosperma* and *Pinus edulis*, in response to changing VPD. Otherwise, we suggest using the following equation to estimate *e*_i_:

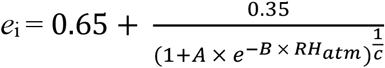

wherein generalized suggested parameters are: A = 2.03, B = 5.179, and C = 0.096, based on fitted RH_cellular_ from the large-scale dataset. More details can be found in the methods section (Equation **14**). The large-scale dataset is a collection of different plant functional groups, and such diversity in the dataset could affect fitted RH_cellular_, perhaps more so at the upper and lower RH_cellular_ limits. For example, variability in stomatal conductance could be responsible for extremely low fitted RH_cellular_ at low RH_atm_, therefore it is likely important to incorporate non-steady state modelling in future studies when RH_atm_ is low. The correction can be refined to specific ecological contexts using site-specific and species-specific information, for example, by calculating a fitted RH_cellular_ based on an existing study of the same species in a similar location, like this study evaluated Scots pine at Hyytiälä. Or, ideally, A and B are refined based on experimental data on species-specific RH_cellular_ responses to RH_atm_ or VPD. Indeed, these are suggested starting points for correcting *e*_i_ for its unsaturation when predicting or reconstructing LWE.

The model optimization estimated that fitted RH_cellular_ for optimal LWE predictions would reach much lower RH_cellular_ than what has been empirically measured by Cernusak et al. (2018) and Wong et al. (2022), especially at low RH_atm_ (Fig. **4**). When the optimization was adjusted to limit fitted RH_cellular_ to measured values, the fitted RH_cellular_ for Hyytiälä was bounded to 80% across all measured RH_atm_ at Hyytiälä, which predicted Δ^18^O_lw_ poorly (Fig. **2G**; Table **2**). At low RH_atm_, the extremely low fitted RH_cellular_ could have been affected by increased stomatal closure, because stomatal closure disrupts the hypothesized relationship between RH_atm_ and RH_cellular_. Also, if Δ^18^O_lw_ was fitted separately to Δ^2^H_lw_ then the fitted RH_cellular_ of Δ^18^O_lw_ is closer to empirical measurements of RH_cellular_ by Cernusak et al. (2018) and Wong et al. (2022), especially at Hyytiälä (Supplemental Fig. **2**). A potential reason for such low fitted RH_cellular_ for Δ^2^H_lw_, is that the CG model did not predict Δ^2^H_lw_ as accurately as Δ^18^O_lw_ in this study (Fig. **2**, **3**; Table **2**, **3**). Therefore, a lower fitted RH_cellular_ for Δ^2^H_lw_ might have been necessary to remediate other sources of error in the CG model for Δ^2^H_lw_. An alternative fitted RH_cellular_ to correct for unsaturated *e*_i_ for Δ^18^O_lw_ only, is based on separate fitting of RH_cellular_ for Δ^18^O_lw_ in the large-scale dataset (A =1088.18, B = 9.81 and C = 3.06, Suppplemental Fig. **2**).

Results suggested that T_leaf_ is important to consider alongside accounting for unsaturated *e*_i_ in future studies, because fitted RH_cellular_ is sensitive to ±2ºC variability in T_leaf_ (Fig. **4**). Cryogenic water extraction artefacts, and xylem sampling effects, may also affect LWE values (Chen *et al*., 2020; Barbeta *et al*., 2022; Diao *et al*., 2022; Nehemy *et al*., 2022).

When applying the Péclet correction, *L* is dependent on assumptions in the CG model (Table **1**; Loucos et al. (2014)). In this study, we tested assumptions that could affect the calculation of *L*. Also, the Péclet and two-pool corrections can affect the accuracy of Δ^2^H_lw_ and Δ^18^O_lw_ predictions differently (Bögelein, Thomas and Kahmen, 2017). Therefore, it is understandable that in this study, the selected Péclet and two-pool corrections were favorable to only a minority of scenarios (Fig. **2**; Table **2**). Importantly, we have shown that it is necessary to develop co-implementation of the two-pool and Péclet with unsaturated *e*_i_ further, because the versions used in this study were incompatible with unsaturated *e*_i_ effects to 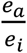 of the CG model (Equation **2;** Fig. **2**; Table **2**). After all, when RH_cellular_ was fitted after the two-pool and Péclet corrections, the fitted RH_cellular_ became unrealistically high (103 – 146% and 105 – 210%, respectively, Supplemental Fig. **1**). It is worthwhile to further explore the co-implementation of two-pool and Péclet corrections alongside adjustments for unsaturated *e*_i_, because Péclet and two-pool corrections have potential to remediate model prediction accuracy bias introduced by unsaturated *e*_i_ via 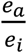 of the CG model (Fig. **2**; Table **2**).

### Unsaturated *e*_i_ effects to fractionation within ε_k_

Results from this study showed that it is not necessary to further explore the effect of unsaturated *e*_i_ to fractionation within ε_k_, by decreased *r* and introduced *r*_i_, for 80 – 100% RH_cellular_, because it had a negligible effect to predicted Δ^2^H_lw_ and Δ^18^O_lw_ (< 0.17‰, Fig. **2 B, C, F, G**). A contributing factor to this finding, is that the influence of ε_k_ to LWE decreases in response to unsaturated *e*_i_, as observed for Hypothesis 1. Therefore, intercellular resistance (*r*_i_) does not need to be incorporated into the calculation of ε_k_ in response to unsaturated *e*_i_, unless the calculation of ε_k_ receives major revision in the future. There is opportunity for future investigations to explore how different calculation techniques for *g*_s_ might be affected by unsaturated *e*_i_ (see Damour *et al*., 2010).

## Conclusion

Our results show that accounting for unsaturated *e*_i_ changes spatiotemporal LWE predictions and can even improve them. We therefore conclude that unsaturated *e*_i_ should be considered as a key modification factor of leaf water stable isotopes in future studies. Particularly, to account for higher influence of Δ_*v*_ and lower influence of ε_*k*_ by decreasing *e*_i_ in 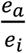 of the CG model (Equation **2**). Corrections which use a constant value of RH_cellular_ when calculating 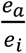 are not effective, likewise it is ineffective to continue to use a constant RH_cellular_ of 100%, when *e*_i_ is assumed to be saturated. We propose a model correction for both Δ^2^H_lw_ and Δ^18^O_lw_ based on RH_atm_, and we suggest that such an approach can alternatively be applied with VPD. This model correction is a starting point for more accurately predicting LWE or reconstructing LWE from plant-derived organic proxies such as tree rings and *n*-alkanes. This may particularly benefit ^2^H interpretations, due to the noticeable improvement on Δ^2^H_lw_ predictions by lowered RH_cellular_ during LWE modelling, but it may not benefit ^18^O interpretations.

## Acknowledgements

Many thanks to Elina Sahlstedt, Giles Young, Kersti Leppä, Eloise Angove, Magdalena Drys, Aleksi Lehtonen, Yann Salmon, Daniel Zannoni and three anonymous reviewers for constructive comments, to Juha Heikkinen for statistical consultation, to Pasi Kolari for providing leaf cuvette gas exchange data, to Aino Seppänen for water extractions and to Daniel Nelson for performing stable isotope measurements.

## Author contributions

YT, PS-A, KTR-G, PK & JB planned, facilitated and/or conducted field work at Hyytiälä. The subsequent manuscript idea was realized by CA, KTR-G, ML & MS. Data processing, isotope modelling, and analyses were conducted by CA and O-PT. CA was responsible for writing the manuscript, with major contributions from KTR-G, ML, MS, YT, AK, O-PT and all authors contributed to the writing of the manuscript.

## Data availability

Review data used in this study is freely available online thanks to Cernusak et al. (2022). The data from Hyytiälä which support findings from this study will be made freely available.

## Supplemental Materials

**Supplemental Table 1** Linear mixed model fits between modelled and observed leaf water deuterium (Δ^2^H_lw_) and oxygen-18 (Δ^18^O_lw_) when intercellular relative humidity (RH_cellular_) has been adjusted in association with, either, enrichment of atmospheric vapor relative to source water (Δ^2^H_wv_ or Δ^18^O_wv_), or, the kinetic fractionation during diffusion through the stomata and boundary layer (ε_k_) in the Craig Gordon model.

**Supplemental Fig. 1** Fitted leaf intercellular space relative humidity, in response to atmospheric relative humidity, with different corrections for leaf water heavy isotope enrichment modelling.

**Supplemental Fig. 2** Fitted leaf intercellular relative humidity, in response to atmospheric relative humidity, separately fitted for each of Δ^18^O_lw_ and Δ^2^H_lw_.

